# Cell-Vision Fusion: A Swin Transformer-based Approach to Predicting Kinase Inhibitor Mechanism of Action from Cell Painting Data

**DOI:** 10.1101/2023.12.13.571534

**Authors:** William Dee, Ines Sequeira, Anna Lobley, Gregory Slabaugh

**Author notes:** Correspondence to: William Dee, Digital Environment Research Institute (DERI), Empire House, 67-75 New Road, London, E1 1HH. Anna Lobley and Gregory Slabaugh should be considered joint senior author.

## Abstract

Image-based profiling of the cellular response to drug compounds has proven to be an effective method to characterize the morphological changes resulting from chemical perturbation experiments. This approach has been useful in the field of drug discovery, ranging from phenotype-based screening to identifying a compound’s mechanism of action or toxicity. As a greater amount of data becomes available however, there are growing demands for deep learning methods to be applied to perturbation data. In this paper we applied the transformer-based SwinV2 computer vision architecture to predict the mechanism of action of 10 kinase inhibitor compounds directly from raw images of the cellular response. This method outperforms the standard approach of using image-based profiles, multidimensional feature set representations generated by bioimaging software. Furthermore, we combined the best performing models for three different data modalities, raw images, image-based profiles and compound chemical structures, to form a fusion model, Cell-Vision Fusion (CVF). This approach classified the kinase inhibitors with 69.79% accuracy and 70.56% F1 score, 4.20% and 5.49% greater, respectively, than the best performing image-based profile method. Our work provides three techniques, specific to Cell Painting images, which enable the SwinV2 architecture to train effectively, and explores approaches to combat the significant batch effects present in large Cell Painting perturbation datasets.

## INTRODUCTION

In the field of drug development, the vastness of the “drug-like” chemical space makes it impractical to explore through clinical trials alone ^1^. As a result, less complex model systems such as cells or tissues are commonly employed for the initial testing of potential drug candidates. Powered by the latest advancements in high-throughput microscopy, image analysis has become a core component for investigating the impact of chemical compounds on cellular morphology, structures and processes ^2^. Image-based profiling (IBP) is one popular method of analyzing the cellular phenotypic changes caused by chemical perturbations. The choice of profiling assay influences the information that can be extracted from the cell’s response, ranging from multi-omic representations to single readouts of cell health, such as viability or apoptosis ^3–5^.

The most commonly applied IBP assay is Cell Painting ^2,6,7^, where six fluorescent dyes are used to stain eight specific cellular components, including the “nucleus, nucleolus, endoplasmic reticulum (ER), Golgi, mitochondria, plasma membrane, cytoplasm, and cytoskeleton” ^2^. A microscope is then used to image the cell’s response across five channels. The standard approach processes these images using a software package, such as CellProfiler ^8^, to extract morphological features at the single-cell level after applying segmentation techniques. These features are aggregated into several thousand profile-level features which aim to represent the overall cellular response to a particular perturbation. The feature sets measure changes to the cell’s size, shape, intensity, protein colocalization and texture ^7,9^. Post-processing steps including normalization, standardization, feature selection and dimensionality reduction are then applied to facilitate downstream supervised or unsupervised machine learning methods ^10^.

Cell Painting has proven effective in the field of drug development, enabling the comparison of cellular response between patients with a disease and healthy patients ^11^, predicting the mitochondrial toxicity of drug compounds ^12^, and for inferring the mechanism of action (MOA) of different compounds ^9,13–15^. The assay captures fundamentally distinct information from transcriptomic or proteomic profiling, while being a higher throughput and lower cost approach to obtaining quantifiable information about a cell ^16–19^.

Despite this, the historical lack of a large-scale dataset, produced using a standardized assay and labelled with ground truth information, has limited the application of deep learning methods in the field. Prior work has typically been restricted in scale, often incorporating fewer than 1,000 perturbations ^13,20–22^, and predominantly utilizing clustering algorithms to differentiate between compounds which often have unrelated mechanisms. This data scarcity, coupled with a growing demand for high throughput approaches to assess cellular phenotypic response ^6,16,20^, prompted the formation of the Joint Undertaking for Morphological Profiling (JUMP) Cell Painting Consortium, which released “cpg0016”, a Cell Painting dataset containing more than 140,000 chemical and genetic perturbations ^16^.

Prior IBP research has postulated whether large Cell Painting datasets, with data produced at multiple centres under varying conditions, could be combined successfully, despite the inevitable batch effects. Novel deep learning solutions are sought-after, given their ability to handle complex, high dimensional data. Therefore, they may be able to model the “subtlety of cellular phenotypes” ^6,20^, while navigating the noise created by confounding factors such as off-target toxicity ^24^. Furthermore, the current standard approach is reliant on single-cell segmentation, making it less relevant for specific cell types, such as adhesion cells which typically grow close together, making their borders difficult to segment ^25^. This limitation has led previous phenotypic profiling research to focus on “broad cell groups, such as lymphocytes, granulocytes, and erythrocytes, that have morphologies easily distinguishable to the human eye” ^10^. Advanced computer vision-based methods can learn directly from the images themselves, rather than being reliant on an intermediate feature set. Firstly, this introduces the possibility of uncovering previously unanticipated biology, not accounted for by the well-established profiling features. Secondly, by learning from the whole image, greater context behind the cellular response may be captured, allowing the model to incorporate a more accurate representation of dynamic spatial features like cell motility. Deep learning-based approaches have proved successful in many biological domains, from the classification of radiographic and pathological images ^26,27^ to industrial-level drug discovery ^11^. They have also recently been applied in the context of image-base profiling to develop phenotypic models of active SARS-CoV-2 infection ^28^ and in the identification of new potential compounds to rejuvenate the immune systems of elderly adults ^29^.

In this paper, we predict the MOA of 10 different kinase inhibitor compounds, tested in the cpg0016 dataset (Figure 1). The MOA of many drugs is often poorly understood at a molecular level, resulting in the high failure rates of clinical drug development ^30^. Phenotypic drug discovery is seen as an alternative to target-based approaches, and one that has led to comparatively high numbers of recent first-in-class molecular entities ^31^. However, determining the MOA of compounds of interest with high accuracy has been a major barrier ^32^, one which may be subsiding with the development of sophisticated phenotypic assays like Cell Painting, in combination with the latest machine learning methods. Imaging offers researchers a bird’s-eye view of cellular phenotypic changes, providing a physical definition of the cellular response. For example, some inhibitors rupture cell membranes or nuclei, others induce gross structural defects or render cells non-functional by causing such significant damage to core machinery to promote senescence. These signals reside in the imaging data and can be used to compare and characterize the multiplicity of effects of different drug compounds.

**Figure 1.**
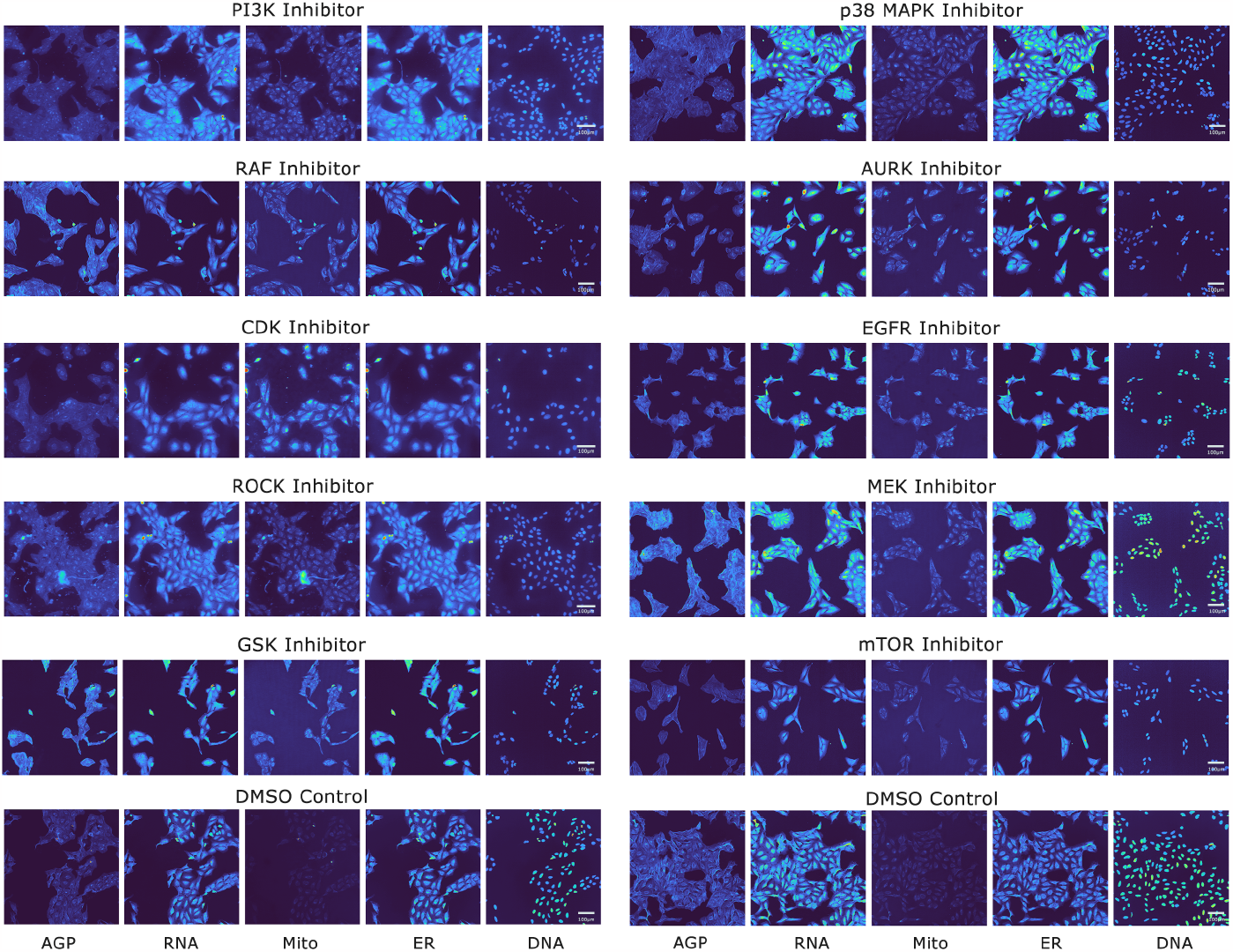
Cell Painted images of cells treated with different kinase inhibitor compounds. Example five-channel Cell Painting images of the 10 kinase inhibitor classes included within the dataset used in this study. The classes include: Phosphoinositide 3-kinase (PI3K), Epidermal growth factor receptor (EGFR), p38 mitogen-activated protein kinase (p38 MAPK), Rapidly Accelerated Fibrosarcoma (RAF), Aurora kinase (AURK), Rho-associated kinases (ROCK), Mitogen-activated protein kinase kinase (MEK), Glycogen synthase kinase (GSK), Cyclin-dependent kinase (CDK) and Mammalian target of rapamycin (mTOR). The five channels (bottom) correspond to the following organelles/cellular compartments: DNA - Nucleus, ER - Endoplasmic reticulum, RNA - Nucleoli, cytoplasmic ribonucleic acid, AGP - F-actin cytoskeleton, Golgi, plasma membrane, Mito - Mitochodria. Dimethyl sulfoxide (DMSO) negative controls are included for direct comparison, but are not classified in this work. In this figure, the PI3K, ROCK and CDK images were produced by source three, DMSO by source 10, and the remaining class images were from experiments performed by source two of the JUMP Consortium.

The kinase inhibitor class was selected for this research due to their importance in the field of drug discovery, particularly in targeted cancer therapy, as well as their well-documented off-target effects which make them, theoretically, a difficult subpopulation to classify with high accuracy ^33,34^. Our class-specific MOA classifier, enabled by the greater data availability present within cpg0016, is therefore distinct from prior MOA research, which has often compared disparate mechanisms such as ATPase inhibitors with retinoid receptor agonists ^15^, or Eg5 inhibitors with microtubule stabilizers ^13^.

Within this work we compare and contrast different computational methods of normalization, standardization, feature selection and data augmentation when training models with the three separate modalities which are present within the cpg0016 dataset – Cell Painting images, image-based profiles, and compound chemical structures. We introduce three methods, specific to five-channel Cell Painting image data, which enable the SwinV2 computer vision architecture ^35^ to classify images directly from the raw pixels with better performance than the standard approach using CellProfiler IBP features. Finally, we combine the three data modalities to form a fusion architecture (Figure 2), referred to as Cell-Vision Fusion (CVF), which outperforms the single modality approaches, confirming similar results found in other Cell Painting papers where more than one modalities were combined ^15,36,37^.

**Figure 2.**
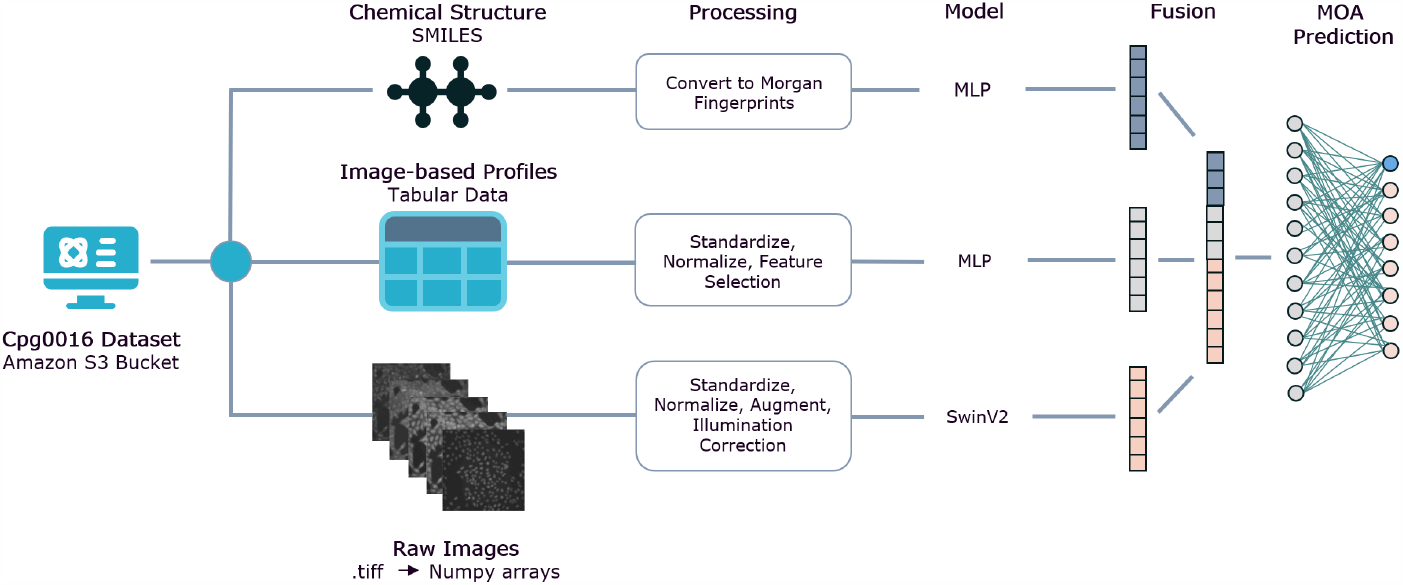
Overview of the Cell-Vision Fusion (CVF) architecture. Specific models for each data modality are each incorporated into the architecture before the outputted embeddings from each are combined using Multi-modal Outer Arithmetic Block (MOAB) fusion ^23^. The fusion model is trained end to end, see Methods for more details.

## RESULTS

### Feature selection

The CellProfiler-produced IBP data contains 4,762 features representing the cellular response to a perturbation in a specific experimental well. To reduce these features to focus on those which are specific to identifying kinase inhibition, we compared the impact on model performance of using a Shapley value ^38^ feature ranking with the current standard approach of using the Pycytominer package (v1.0.1) ^39^. The Shapley importance values revealed that there were a relatively small number of important features. The top 20 features had substantially higher importance than the other features, and there were many features in the long tail of the distribution with very little positive contribution to model performance (Figure 3A). This observation was confirmed when these ranked features were used to select model features. Figure 3B shows the cross-validated accuracy and F1 scores of an XGBoost model, trained with varying numbers of features, revealing that, past the top 150 Shapley ranked features, including more features reduces performance. 150 features were selected as the optimal number due to the lower standard deviation between cross-validation folds (the shaded areas in Figure 3B), as well as being the number which elicited the highest F1 score. This number of features is similar to the 184 Cell Painting features used by Seal et al. ^37^ to predict assay hit calls.

**Figure 3.**
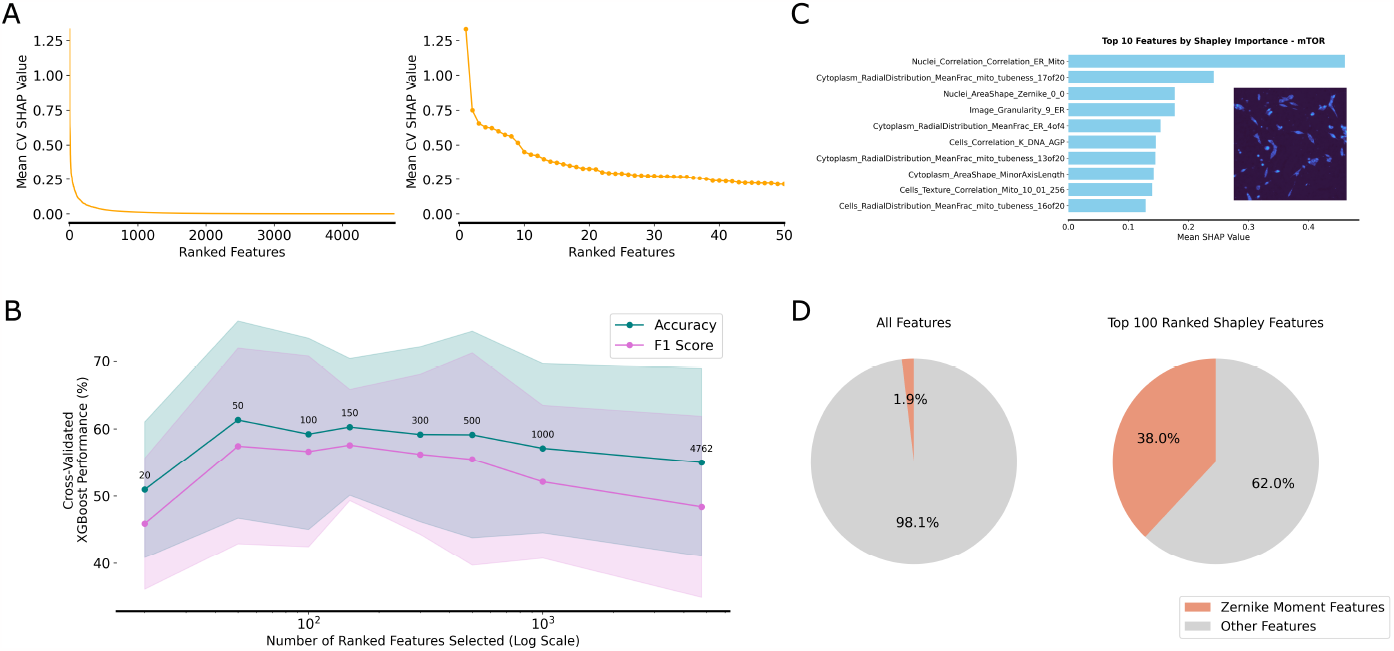
Displaying the importance and intepretability of Shapley-ranked CellProfiler feature rankings. A) Shapley-ranked feature importance plot showing two different views of the same results, where all features are shown (left) and where the top 50 features are plotted (right). This displays the drop-off in feature importance past the top 20 Shapley-ranked features when measuring their importance for XGBoost model prediction. B) An illustration of the performance of an XGBoost model (before hyperparameter tuning) which incorporates the top n-ranked features selected according Shapley value. Features are plotted using a log-scale to enable comparison. The shaded area represents the range between accuracy and F1 scores between five cross-validation folds. C) An example Shapley value importance table for the top 10 ranked features for predicting the mTOR inhibitor class, alongside an example image from the mitochondria channel of a mTOR inhibitor datapoint. D) A pie chart showing the importance of Zernike moment features in the Shapley value rankings, accounting for 1.9% of the whole CellProfiler feature set, but 38% of the top 100 Shapley-ranked features.

Comparatively, applying Pycytominer’s feature selection module resulted in 871 retained features. Using these features in the same baseline XGBoost model achieved an average cross-validated accuracy of 57.05% and F1 score of 53.43%, underperforming the 150 Shapley feature model which recorded 60.26% accuracy and 57.57% F1 score. In the following sections we compare how these two feature selection methods perform when combined with different normalization and standardization approaches, as well as when using a range of different model types.

The Shapley value analysis also highlighted two important results. Firstly, prior to performing quality control (see Methods: Quality control), the feature with the greatest importance was one representing the image quality of the DNA channel, implying image quality was impacting model performance to a greater degree than the underlying biological signal. Post quality control measures, these image quality features dropped significantly in terms of Shapley importance, with the DNA image quality feature falling to the 110th rank. Secondly, the importance of Zernike moment ^40^ features was highlighted. Zernike moments are a set of orthogonal complex-valued polynomials defined over a unit disk which are widely used in image processing and computer vision for shape analysis and pattern recognition. In this context they have been applied by CellProfiler to represent the shape of the single cells captured in the images. Zernike moment features account for only 1.88% of the original 4,762 feature set, but 38% of the top 100 Shapley-ranked features (Figure 3D), displaying the significance of cellular morphology shape changes when classifying kinase inhibitors. This finding aligns with prior kinase inhibitor research, which shows cell shape can be significantly altered by inhibitors preventing kinase proteins from regulating cytoskeleton dynamics ^41,42^, as well as affecting cell adhesion, migration and death via apoptosis ^5,43–45^. This also suggests computer vision models, given their aptitude for object detection, are well-suited for the task of differentiating between cellular response where morphological changes are so prominent.

One further strength of using Shapley values is the ability to be able to interpret their individual impact on the model for each separate class. Figure 3C shows the Shapley importance values for the top 10 features that contributed to the classification of the mTOR class. The top ranking feature for mTOR relates to the correlation between the nuclei and the endoplasmic reticulum and mitochondria channels. mTOR is an important regulator of mitochondrial function ^46,47^ and plays a regulatory role in endoplasmic reticulum stress ^48^ where the inhibition of mTORC1 signalling has been found to attenuate endoplasmic reticulum stress-induced apoptosis ^49^. Furthermore, there are several features representing “tubeness”, i.e., how “tube-like” the mitochondrial structures in the images are, as well as their distribution or prevalence throughout the image. This feature of mTOR inhibition can be observed in the example mitochondrial channel image in Figure 3C.

The Shapley feature ranking can therefore be useful to provide an understanding of how the CellProfiler features relate to underlying morphological changes as well as being directly relatable to literature. Additionally, these rankings may provide new insights into mechanisms of action, which can form the basis for experimental studies or applications. The top 10 Shapley-ranked CellProfiler features for each class can be found in Supplemental Note S1.

### Reducing batch effects: Image-based profiles

Batch effects are a key consideration when training models on Cell Painting data ^2,3,6,9^, especially in the case of cpg0016 where experiments were performed by different contributing partners (referred to as “sources”), under a variety of experimental conditions (see Methods: Data acquisition). To attempt to reduce the impact of these batch effects, different normalization and/or standardization methods were employed. Normalization scales and centres data within a specific range (often between zero and one) and is commonly used when algorithms require input values in these ranges to function effectively. Standardization transforms data to have zero mean and unit variance, centering the data around zero. Prior work has shown using control samples from the same experimental plate to normalize/standardize data can help to partially eliminate batch effects ^2,3,21^. To visualize the impact of these effects within our kinase inhibitor dataset, and to gain some insight into how well each approach counteracted them, while retaining the underlying biological signal, we used Uniform Manifold Approximation and Projection (UMAP) ^50^.

UMAP plots were created showing how the IBP datapoints clustered in two dimensions. Three different labels were used to colour the datapoints, being the microscope which was used to take the image (Figure 4A), the source where the experiment was performed (Figure 4B) and the MOA of each compound (Figure 4C). Each plot is titled according to the processing techniques which were applied, where “Baseline” reflects the raw data prior to these steps. The baseline plots show the extreme impact of the batch effects in the data, causing datapoints to cluster according to the microscope and source, but showing no association between compounds with the same MOA.

**Figure 4.**
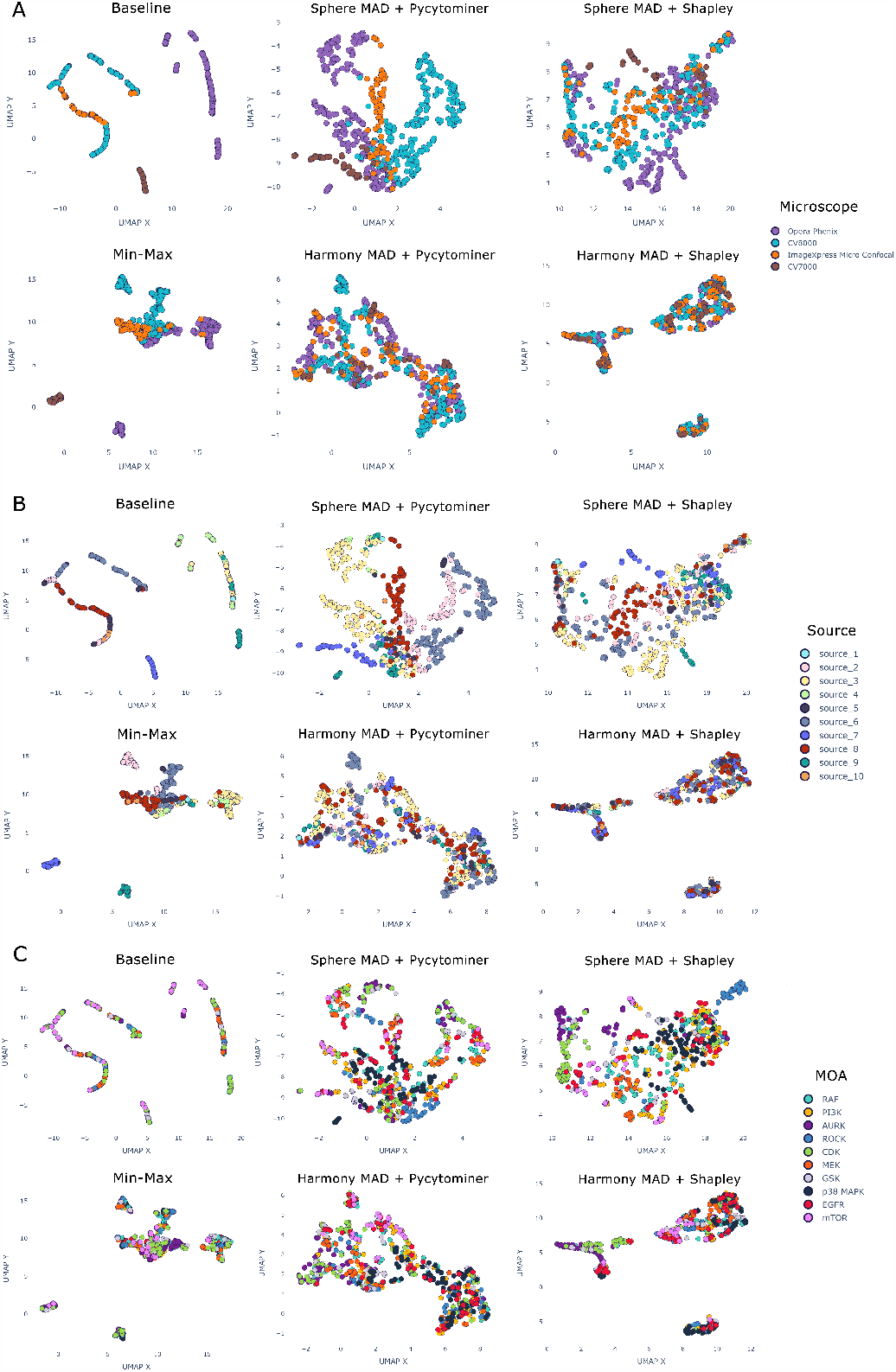
Reducing the impact of batch effects by normalizing and standardizing the image-based profile data. UMAP plots displaying the IBP data, reduced to two dimensions. The data is coloured according to three different labels: A) the microscope used to take the five-channel Cell Painting image, B) the contributing “source” (i.e. JUMP partner) where the experiment was carried out, and C) the compound’s mechanism of action. The plots show the image datapoints before any normalization or standardization has been applied (“Baseline”), after min-max normalization, and by applying combinations of either spherization or Harmony in combination with either Pycytominer or Shapley-ranking based feature selection. An ideal plot would display no clustering according to the microscope type in A or originating source in B (as these are not biologically relevant), and instead would show datapoints clustering close together if the compounds they represent operate with the same mechanism (C), and thus should result in similar observed phenotypic changes.

Using median absolute deviation (MAD) normalization in combination with either spherization or Harmony (see Methods for more detail) resulted in a clearer separation of the datapoints compared to the baseline approach or min-max normalization. While Harmony appeared to be more adept at reducing the impact of the microscope, overall, the datapoint MOA clusters are less well aligned than when using MAD normalization, spherization and Shapley-ranked feature selection (“Sphere MAD + Shapley”). Spherization and MAD normalization works more effectively when using only the top 150 Shapley-ranked features, rather than the 871 features retained by Pycytominer. This suggests some of Pycytominer features are highly correlated to the underlying microscope and source batch effects, and removing them would improve a model’s ability to classify by MOA. Based on these plots we would expect a classifier, trained on the Shapley-reduced feature set, MAD normalized and using a spherization transformation to have the best performance (see Classifying Kinase Inhibitor MOA). Looking at individual MOA classes (Figure 4C) we can observe that Rho-associated kinase (ROCK), Aurora kinase (AURK) and Cyclin-dependent kinase (CDK) inhibitors appear most well-distinguished. In contrast, Rapidly Accelerated Fibrosarcoma (RAF), Epidermal growth factor receptor (EGFR) and Phosphoinositide 3kinase (PI3K) appear the least well clustered, implying these classes will be the most difficult to classify with high accuracy.

### Reducing batch effects: Raw images

UMAP embeddings were also plotted for the raw image data, before and after using per-channel normalization (PCN) and plate-wise, channel-wise standardization (PCS) (Figure 5). PCN min-max normalizes the image channels separately between zero and one, while PCS standardizes each channel based on either the DMSO negative controls or the positive control compound average pixel values for the specific plate where the experiment image was captured (see Methods: Raw Images: Normalization and standardization). Both methods perform similarly in terms of counteracting the obscuring batch effects visible in the baseline UMAP embeddings, caused by both the experiment source and microscope used. Unlike the UMAP plots in Figure 4, where the ROCK, AURK and CDK inhibtors partially clustered according to MOA, there are no clear MOA clusters revealed by the UMAP embeddings of the images. This was anticipated given these embeddings were obtained by passing each image through a pre-trained ResNet-50 classifier that had not been fine-tuned on any Cell Painting data, and because the information contained in the raw pixels is likely too complex to represent effectively in two dimensions. A computer vision architecture, specifically trained on Cell Painting data, is therefore likely required to separate classes effectively. The positive impact these methods had on eliminating batch effects can be evidenced in the Table 1 results and Figure 6F, which show the model performance improvements when PCN and PCS are applied, in addition to using channel-weighted data augmentation (CWA) throughout training (see Methods: Raw images: Data augmentation).

**Table 1.** Five-fold cross-validated results for predicting kinase inhibitor mechanism of action. Results showing the performance of different models when using different data modalities - raw Cell Painting images, image-based profiles and the compound chemical structures, converted to Morgan fingerprints. The Fusion model incorporates all three modalities and fuses the MLP models for both compound structure and IBP data with the SwinV2 model for the images.

| Approach | Accuracy | F1 Score (Macro) | Precision | Recall | AUPR (Macro) | AUC (Macro) |
| --- | --- | --- | --- | --- | --- | --- |
| **Image-based profiles** |  |  |  |  |  |  |
| **XGBoost** | 65.59 | 63.99 | 66.12 | 64.32 | 68.61 | 89.02 |
| **MLP** | 65.59 | 65.07 | 68.07 | 64.55 | 67.63 | 89.45 |
| **Raw images** |  |  |  |  |  |  |
| **SwinV2** | 66.67 | 69.59 | 72.24 | 68.92 | 67.43 | 89.04 |
| **EfficientNet** | 62.05 | 54.17 | 52.67 | 61.66 | 65.40 | 80.30 |
| **Compound structure** |  |  |  |  |  |  |
| **XGBoost** | 55.21 | 48.45 | 49.58 | 49.68 | 55.90 | 83.25 |
| **MLP** | 58.33 | 56.08 | 58.19 | 55.94 | 60.31 | 84.34 |
| **Combined data modalities** |  |  |  |  |  |  |
| **Cell-Vision Fusion** | 69.79 | 70.56 | 73.40 | 69.87 | 74.12 | 90.73 |

**Figure 5.**
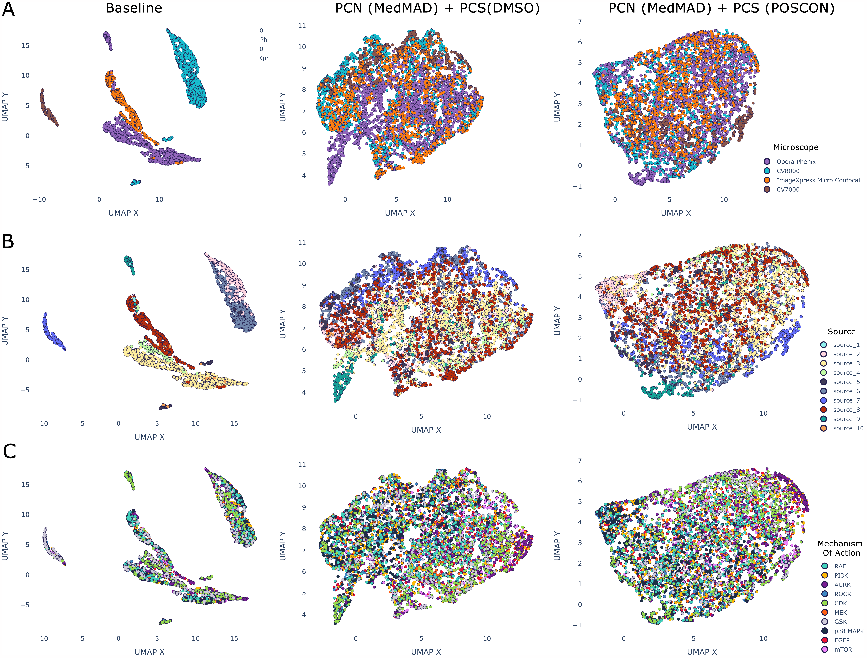
Reducing the impact of batch effects by normalizing and standardizing the image data. UMAP plots displaying the raw image data, reduced to two dimensions after being passed through a ResNet-50 classifier to obtain embeddings. The data is coloured according to three different labels: A) the microscope used to take the five-channel Cell Painting image, B) the contributing “source” (i.e. JUMP partner) where the experiment was carried out, and C) the compound’s mechanism of action. The plots show the image datapoints before any normalization or standardization has been applied (“Baseline”), and after PCN and MedMAD PCS (see Methods: Raw images: Normalization and Standardization for more details) has been performed, using either the responses of the DMSO negative or positive control (POSCON) wells to reduce batch effects.

**Figure 6.**
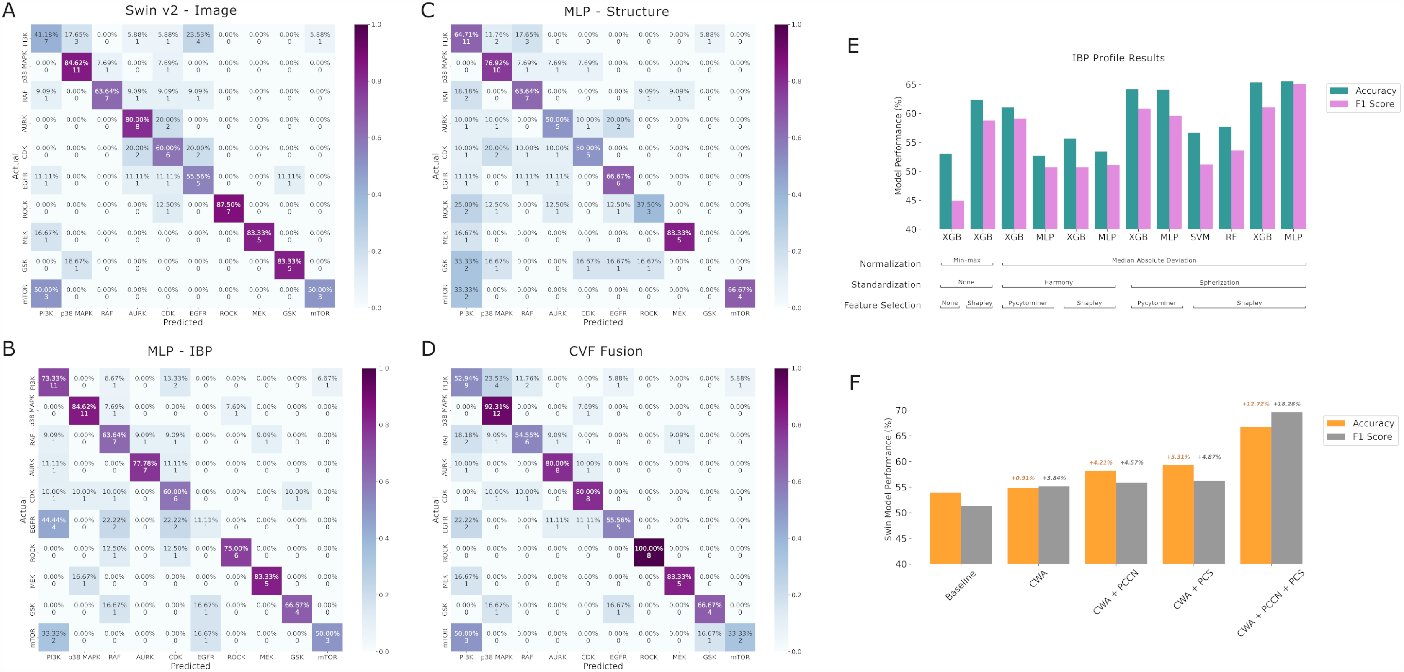
Model predictive performance comparing different model types and processing steps. A-D) Confusion matrices showing the model predictions compared with the actual compound labels for the A) SwinV2 model, trained on the raw images, B) the MLP model, trained on the IBP data, C) the MLP model, trained on the compound structure-based Morgan Fingerprints and D) the CVF fusion model, combining the models in A), B) and C). E) shows the impact of different normalization, standardization, feature selection and model archictectures when predicting MOA using the IBP data. F) shows the comparative performance of the SwinV2 method when utilizing the CWA, PCN and PCS techniques over a baseline approach where none are applied.

### Classifying Kinase Inhibitor MOA

Five-fold average test set performance metrics were computed, comparing models trained using the three different data modalities (image-based profiles, images and compound structure), as well as the fusion model architecture (Table 1).

The best performing model trained on a single modality was the SwinV2 model as it achieved greater average accuracy, F1 score, precision, recall and AUPR metrics than all other single modality approaches. Despite also being trained on images, EfficientNet performed comparatively worse, achieving accuracy and F1 scores of 62.05% and 54.17% respectively, compared to SwinV2’s 66.67% and 69.59%, reflecting EfficientNet’s inability to classify the EGFR or mTOR inhibitor classes. For both the IBP and compound structure modalities, the MLP models sourced from prior work ^3,15^ outperformed the less complex XGBoost approach. However, the improvement was only marginal for the IBP data, likely due to the XGBoost-based Shapley feature selection.

The confusion matrices of the best performing models for each modality enable a comparison of how well suited each modality is to predicting each MOA class (Figure 6 A-D). The SwinV2 model is comparatively worse at predicting the majority class (PI3K inhibitors), the IBP-based model predicts the EGFR class with only 11.11% accuracy, while the compound structure-based MLP finds it most difficult to predict the ROCK and GSK classes. The only consistently well-predicted class is p38 MAPK.

Overall the fusion model achieved the best performance, gaining a macro-averaged F1 score of 70.56% alongside strong performance in the other measure metrics. This demonstrates there was a clear synergistic impact from combining the different modalities and each model architecture was extracting partially unique information from the data. This was suggested by the confusion matrices which show distinctly different predictions per class between the approaches. The performance improvements gained when applying different model architectures, normalization, standardization and feature selection methods were compared for the image-based profile data (Figure 6E) and the raw image data (Figure 6F). The best performing IBP-based MLP and XGBoost models (results shown in Table 1) both utilized MAD normalization, spherization and the top 150 Shapley-ranked features, as predicted from observing the UMAP embedding plots (Figure 4).

For the image data, channel-weighted augmentation (CWA) boosted performance by 0.91% and 3.84% over the respective accuracy and F1 score of the baseline approach (Figure 6F). Combining per-channel normalization (PCN) with CWA improved accuracy by 4.21% and F1 score by 4.57%, while plate-wise, channel-wise standardization (PCS) in combination with CWA increased accuracy and F1 score by 5.31% and 4.87%. However, using all three techniques in unison resulted in a much greater performance improvement, increasing accuracy by 12.72% and F1 score by 18.28%, ultimately enabling the SwinV2 architecture to outperform the IBP-based approaches.

## DISCUSSION

We have successfully applied the SwinV2 computer vision architecture to predict kinase inhibitor MOA directly from Cell Painting images with performance exceeding that of the current standard IBP-based approach. Our work confirms assumptions that additional context can be derived by incorporating the whole images into a deep learning model ^16^, providing additional information that is not captured by the CellProfiler features derived from single-cell segmentations. Combining three separate data modalities into the Cell-Vision Fusion architecture proved to be beneficial for model performance overall. When put in the context of a compound screen of thousands of potential drug candidates, the 4.2% improvement in both accuracy and precision over the best image-based profile approach is impactful in terms of correctly identified MOAs which can inform further testing.

Our work has also shown the batch effects present in cpg0016 can be at least partially eliminated, through various normalization and standardization techniques. In the case of using the images, taking a channel-based approach to augmentation, as well as normalization and standardization, proved to be beneficially additive to the model during training, reflecting the channel-specific information included in the Cell Painting five-channel format. Our work also explored the benefits of using Shapley rankings to perform feature selection over the current standard approach to use Pycytominer. While the performance gain was not excessive (1.3% greater accuracy and 3.5% greater F1 score - Figure 6E), the additional interpretability of the model features with greatest importance for class-specific prediction is a noteworthy benefit as it provides comparability with known literature, as well as the possibility of uncovering previously unanticipated biology.

ROCK, AURK, p38 MAPK and MEK inhibitor classes are well predicted across both image and IBP modalities. All of these MOA classes were shown to cluster more distinctly than other classes in the UMAP diagrams (Figure 4C and Figure 5C). Furthermore, the Shapley importance features tend to be more unique for these classes, suggesting there are definitive morphological changes specific to the inhibition of those proteins which are captured by CellProfiler and are well represented in the amalgamated profiles. Lastly, the DepMap TPM gene expression values (see Supplemental Note S2) for genes related to these inhibited proteins are relatively high compared to other classes in our dataset, suggesting their inhibition results in a strong phenotypic response which is more easily able to be captured by morphological feature representations.

Looking at the individual MOA class predictions across model types, there are some insights into why particular inhibitors may have been classified with lower comparative accuracy. The SwinV2 model exhibits a lower predictive accuracy on the majority PI3K class (17.7% of all compounds) with 41.18% accuracy, compared to 73.33% using the IBP data and 64.71% with the compound structures. The image-based model commonly miss-classifies PI3K as EGFR, while the IBP-based model does the opposite, identifying EGFR samples with only 11.11% accuracy. This may reflect the fact the PI3K/Akt pathway is one of the major signalling pathways downstream from EGFR ^51,52^, and PI3K/Akt signal transduction has been shown to be inhibited by antibodies binding directly to EGFRs ^53^. On inspection of the highly ranked Shapley features for both classes (see Supplemental Note S1), both classes share common important features relating to the F-actin cytoskeleton, Golgi and plasma membrane (AGP) channel. The EGFR is an actin-binding protein ^54^ which has been shown to reorganize the junctional actin cytoskeleton ^55^. PI3K can also disrupt the actin cytoskeleton ^56^, while both contribute to actin filament remodelling throughout the cell ^57,58^. EGFR is also highly connected to the functioning of the Golgi apparatus in protein transport ^59,60^ and is a plasma membrane receptor. Comparatively, PI3K-generated PIP3 is localized to the plasma membrane and is involved in protein kinase localization and activation as well as the EGFR regulation of membrane ruffling ^61^. These impacts to both cytoskeletal rearrangement, membrane activity and cell motility are likely causes for the models to misclassify PI3K and EGFR when only the cellular phenotypic response is considered. Notably, the prediction of both classes improves when only relying on compound structure. Futhermore, both EGFR and PI3K-related DepMap TPM gene expression values were among the lowest of all the kinase inhibitor classes tested within cpg0016 (see Supplemental Note S2), implying there would be a less significant phenotypic response to these compounds in U2OS cells specifically. For similar inter-related pathway reasons, mTOR, which operates downstream from PI3K/Akt, is most commonly misclassified as PI3K by all models. Additionally, both classes exhibit phenotypic shape changes that elongate the mitochondria, demonstrated in the images, as well as the high ranking of “tubeness” features for both classes in the Shapley values. This issue is likely exacerbated by the fact that there are only six unique mTOR inhibitors in the dataset, compared to 17 PI3K inhibitors.

EfficientNet comparatively performed poorly at this prediction task. There are several factors which may have contributed to this result. Firstly, the reduction in image size to 240 × 240 pixels compared with the 896 × 896-pixel images fed into the SwinV2 model, may have meant EfficientNet missed some of the fine-grained contextual information retained at higher resolutions. Also, due to being convolution-based, EfficientNet is better suited to modelling local contextual interactions, rather than the global context captured by transformer-based models owing to the attention mechanism. EfficientNet would likely be more adept at modelling single-cell or smaller image crops, compared to being passed the whole image. However, this approach would likely yield similar information to the CellProfiler-generated IBP data, therefore negating the necessity to use the images directly. The results do align with the work of Tian et al. ^15^, where an EfficientNet model achieved an overall macro-average F1 score of 81%, but only 48% on the AURK inhibitor class, despite AURK being classified against more disparate classes than it was in this work.

One of the major limitations in the study is the use of a single cell line – U2OS. It is possible, therefore, that models trained on this dataset would fail to generalize with similar performance to a different cancer, or non-cancer, cell line. Similarly, experiments were assayed at the same concentration (10uM, excluding Source seven – see Experimental Conditions). It is highly likely that different compounds have varying potencies and incorporating IC50 information for each compound, or performing experiments at multiple doses could result in more accurate identification of activity. Some evidence of this can be seen in Figure 4B and Figure 5B, where datapoints in Source 7 still appear to cluster according to source post-normalization. Our approach also excluded compounds with multiple target classes (i.e., PI3K/mTOR inhibitors), focusing on compounds with a single documented target. This simplistic approach is not reflective of how the majority of kinase inhibitors function, especially when “polypharmacology is the rule rather than the exception for small molecules” ^21^.

Additionally, the computational complexity in terms of compute resources, energy cost and expertise needed to design and implement the fusion architecture compared with an XG-Boost model trained on IBP data is vastly more demanding. This work, however, sets an initial benchmark for what can be achieved by leveraging computer vision models in this setting. While over 5,000 images were used to train/validate the SwinV2 model, the actual number of unique kinase inhibitor compounds was only 96, revealing a lack of diversity in the data. Utilizing the methods put forward in this paper did allow the computer vision architecture to learn effectively, however this constrained data situation is not typically where deep learning-based modern computer vision methods thrive. A larger, more diverse dataset could arguably yield greater performance gains over the more traditional approaches to classifying Cell Painting data. For further work it would also be interesting to evaluate the impact of including proteomic or transcriptomic feature sets as additional data modalities, to compare and contrast their relative strengths on a class-specific basis.

## METHODS

### Data acquisition

The cpg0016 dataset is part of the JUMP Cell Painting collection of datasets which were either produced or reprocessed by the JUMP Consortium ^16^. Cpg0016 is the largest dataset in the collection, containing cell painting data reflecting 116,753 chemical compound perturbations, as well as 23,119 different genetic perturbations. The experiments which produced this data were conducted across 11 different partner sites (referred to as “sources”) where they were assayed using different instruments and microscopes as detailed in Chandrasekaran et al. ^16^

In this work we have curated a subset dataset (see Data selection) from the compound perturbations within cpg0016. The experimental conditions which the JUMP Consortium used to produce these perturbations are summarised below:

- **Cell type:** The consortium compared both the U2OS and the A549 cell lines in the pilot cpg000 experiment, eventually deciding to use U2OS epithelial cells for cpg0016 as the experimental responses were similarly robust but there has been more prior work performed in this field with U2OS cells. U2OS cells are tractable in the lab because of their high replication rate and relative stability in experimental systems. This gives the assays more chance of being reproducible and thus makes U2OS an obvious choice for large-scale perturbation experiments.
- **Time point:** Different time points were also assessed in pilot experiments, resulting in 48 hours being selected for compound perturbations. This was decided on the basis of observed effectiveness of the treatment within the time period, combined with a lack of cell death.
- **Reagent vendor:** PerkinElmer provided the PhenoVue Cell Painting Kit 2.0 to the Consortium.
- **Microtiter plates:** PerkinElmer Cell Carrier Ultra plates were selected on the basis that they minimized solution evaporation in the outermost wells. Two sources used 1,536-well plates (source one and nine), while the rest used 384-well plates.
- **Fields of view:** Within each well, multiple images of the cellular response were taken, referred to as “fields of view”. The number of fields varied according to source; with source two and 10 capturing six, sources three through eight, 11 and 15 imaging nine, and sources one and nine capturing four.
- **Compound concentration:** Treatment compounds were assayed at 10uM by all sources, excluding source seven, which was administered at 0.625uM to provide some comparative variation. Positive control compounds were assayed at 5uM.
- **Replicates:** Members of the consortium shared compounds between each other during the data gathering process. This ensured at least five replicates of each compound were collected across five different source sites. Some compounds, such as those included within the positive control plates/ wells were replicated across each batch, and as such have many more replicates.
- **Microscopes:** Five different types of microscopes were used – the PerkinElmer Opera Phenix, ImageXpress Micro confocal, Yokogawa CV8000, Yokogawa CV7000, and PerkinElmer Operetta (not used for any of the images in this study). Lenses were either set to widefield or confocal mode. Exact settings can be found within the supplementary Tables 4 and 6 of Chandrasekaran et al. ^16^’s report.
- **Cell painting assay:** The Cell Painting v3 protocol was used to stain and fluoresce the following organelles across five channels: mitochondria (Mito), nucleus (DNA), nucleoli and cytoplasmic RNA (RNA), endoplasmic reticulum (ER), Golgi and plasma membrane and the actin cytoskeleton (AGP) ^7^.
- **Control wells:** Within each plate, the outermost wells are used for positive control compounds, while the next outer column of wells is used to host DMSO controls. DMSO is a frequently used solvent in drug discovery as it has the ability to dissolve a wide range of substances ^62^. Since the solvent is present in each compound treatment, DMSO-only wells are often used as controls to isolate the impact of the solvent itself.

The cpg0016 dataset consists of two core components associated with each perturbation (as of December 2023) – raw images and CellProfiler ^8^ image-based profiles. Both components can be accessed via the [cpg0016 Amazon s3 storage bucket](https://registry.opendata.aws/cellpainting-gallery/).

The raw images are stored in the .tiff file format, with image dimensions ranging from 970 × 970 pixels (source six) to 1,280 × 1,080 (source seven). The image specifications by source are listed in Supplemental Note S3. The JUMPCP Consortium used the CellProfiler software ^8^ to process the raw images into aggregated profiles. The pipeline is described in detail in Chandrasekaran et al. ^16^, but is summarized below.

The s3 bucket contains illumination functions which can be used to correct for the intensity of light imparted unevenly by the microscope, resulting in vignetting. The function is provided as an array, equal in size to the underlying Cell Painting image, which corresponds to illumination intensity per pixel. When the raw image is divided by this function, the impact of the microscope’s illumination is reduced.

CellProfiler contains a suite of segmentation methods which can be used for identifying single cells from raw images. The segmentation approaches chosen by the JUMP Consortium are set out in the [Broad Institute’s imaging production pipeline](https://github.com/broadinstitute/imaging-platform-pipelines/tree/master/JUMP_production). Within each image, each single cell is segmented before features, spanning seven different feature groups, are extracted. Pycytominer is then used to aggregate the singlecell features to create well-level consensus profiles, referred to as image-based profiles.

### Quality control

As of December 2023, quality control results have not been released for the cpg0016 dataset. Therefore, measures were established for the removal of low-quality images, such as those with high levels of blur or saturation, or images with various artefacts present.

Included within the image-based profile data for each well are CellProfiler features which represent image quality metrics. These features are described in full in the [CellProfiler manual v4.0.5](https://cellprofiler-manual.s3.amazonaws.com/CellProfiler-4.0.5/modules/measurement.html). All features relating to image blur, saturation or focus were extracted for the kinase inhibitor dataset and min-max normalized. The saturation scores were heavily right skewed and so were transformed to a normal distribution using sklearn’s “QuantileTransformer” module. to align with the blur and focus score metrics. On inspection of the images, it was deemed that images with a high or low level of saturation were of poor quality. A tanh transformation was then used to shift all high or low saturation images to be high scoring. The focus scores were inverted to ensure that highly scoring images were low focus, rather than high. Datapoints falling above the 90th percentile in any of these categories were then excluded from selection, this reduced the available data by 24.6%, from 18,503 samples to 13,951. Examples of low-quality images, as well as further details about the quality control process used in this study can be found in Supplemental Note S4 as well as the associated GitHub repository for this work.

### Data selection

The compound perturbations within cpg0016 are not labelled with MOA. Therefore, instructions were followed from the [JUMP Consortium’s Compound Annotator repository](https://github.com/jump-cellpainting/compound-annotator), to annotate the perturbations with MOA labels from both the [Drug Repurposing Hub’s v3/24/2020](https://repo-hub.broadinstitute.org/repurposing#download-data)^63^ annotations, as well as [ChEMBL’s drug mechanisms data portal v33](https://www.ebi.ac.uk/chembl/g/#browse/mechanisms_of_action).

From this list of compounds matched with an MOA, kinase inhibitor compounds were selected. We assigned the MOA labels based on the main target of the inhibitor, i.e., a compound targetting either PIK3CA or PIK3CG would be labelled as PI3K, although compounds with multi-class targets, for example a compound targetting both mTOR and PI3CA, were excluded. Since the cpg0016 dataset contains only the U2OS cell type, a separate literature review was conducted to identify which kinase inhibitors would likely elicit an observable phenotypic impact on the U2OS cells specifically. Kinase inhibitors which did not fulfil this criterion were subsequently filtered out from selection, since it would be futile to try to differentiate between compounds with minimal phenotypic impact. This process is outlined in detail in Supplemental Note S2.

After quality control was performed on the remaining datapoints, samples were selected according to the availability of replicates. Compounds with a minimum of four replicates were chosen, limiting each unique compound to a ten-replicate maximum in the dataset, to ensure there wasn’t a severe imbalance between compounds. A non-static replicate number was chosen to investigate the impact on model predictive ability caused by the presence of replicates.

The resulting dataset is displayed in Table 2. Exact compound details, along with cross-validation splits, can be found in the GitHub repository for this work.

**Table 2.** Kinase inhibitor dataset overview. A table displaying the different kinase inhibitors included within the dataset, including the number of unqiue compounds for each class, how many different experimental wells these selected compounds relate to, and the number of Cell Painting images associated with those wells. † Number of well datapoints = unique compounds * number of replicates ‡ Number of images = number of well datapoints * number of fields of view

| Mechanism of Action | Unique Compounds | Well Datapoints<sup>†</sup> | Number of Images<sup>‡</sup> |
| --- | --- | --- | --- |
| PI3K | 17 | 102 | 838 |
| p38 MAPK | 13 | 76 | 595 |
| RAF | 11 | 75 | 607 |
| AURK | 10 | 70 | 566 |
| CDK | 10 | 74 | 586 |
| EGFR | 9 | 67 | 558 |
| ROCK | 8 | 57 | 463 |
| MEK | 6 | 41 | 345 |
| GSK | 6 | 40 | 348 |
| mTOR | 6 | 33 | 285 |
| **Total** | **96** | **635** | **5,191** |

### Image-based profiles: Feature selection

In this paper we compared two different approaches to feature selection. The standard process that has been applied in prior work ^3,6^ involves removing any features: with missing data, low variance (typically setting the threshold to 1.0), which are highly correlated (using a Pearson correlation coefficient score of 0.9 as the cutoff), and any “block-list” features which have been previously identified as “unstable or noisy” ^9^. The full list of blocklisted features can be found at [CellProfiler Blocklist v3](https://figshare.com/articles/dataset/Blacklist_Features_-_Cell_Profiler/10255811/3). The pycytominer package was utilized for this feature selection approach, using the “drop_na_columns”, “variance_threshold”, “correlation_threshold” and “blocklist” functions.

In addition to the standard approach, we also compared model performance when reducing the feature set according to each feature’s Shapley importance values ^38^. Shapley values are a concept originally taken from cooperative game theory, in the context of machine learning they can be used to assess the relative contribution of each feature towards the eventual classification. To calculate feature importance, models are created with all possible combinations of feature sets, observing the impact of feature exclusion or inclusion on a model’s performance. Over the many permutations a value can be extracted, representing the features’ overall impact. This Shapley value can then be used to rank features.

Tree-based models have an inherent structure which allows Shapley values to be calculated efficiently as predictions are made by aggregating combinations of features in a hierarchical manner. XGBoost was used to generate Shapley values for the kinase inhibitor dataset, using the training data to determine feature importance for each cross-validation fold. To determine the number of features to select, the Shapley values were plotted and the “elbow method” was used to visually observe where the marginal contribution from additional feature inclusion was negligible. To aid interpretability of this method, the contribution of each feature towards each predicted class was plotted for the top 10 features.

### Image-based profiles: Normalization and standardization

Several different normalization and standardization approaches were tested with the dataset, post-feature selection. These approaches were applied in an effort to reduce the impact of batch effects. Currently no consistently reliable approach for the batch correction of image-based profiles has been identified, although Arevalo et al. ^2^ provided a systematic evaluation of seven different methods.

Arevalo et al. ^2^ found Harmony ^64^, outperformed the other methods consistently across a range of scenarios. This held true for “Scenario 4” which involved batch correction for a dataset including multiple microscope types and laboratories, few compounds, and multiple replicates – similar to the problem faced in this paper.

Harmony was originally designed for single-cell RNA sequencing batch correction. It starts by using a low-dimensional cell embedding, i.e., principal components analysis (PCA) ^65^, before grouping cells into multi-dataset clusters using soft clustering, and then computing cluster-specific correction factors based on the centroids of these clusters. The resulting clusters correspond to individual cell types and states, resulting in correction factors which allow Harmony to learn a linear adjustment function that is sensitive to cellular phenotypes. The harmonization process iterates until cell cluster assignments stabilize.

We compared this method to the “Baseline” approach in Arevalo et al. ^2^’s work, being Median Absolute Deviation (MAD) normalization plus a spherize transformation, which has proved effective in prior research ^3,11,66^. The “mad_robustize” function from the pycytominer package was used for the normalization, scaling features individually by subtracting the median and dividing by the median absolute deviation of the DMSO control sample features found on the same plate. Spherization was applied to counteract the impact of well positioning ^67,68^. Similar to Way et al. ^3^‘s approach, we used the zero-phase whitening filter (ZCA) applied to the profile correlation matrix to “minimize the absolute difference between the transformed and untransformed profiles” ^69^. Mix-max normalization was also tested as a relatively more basic normalization approach to provide context for the impact of the other methods. Arevalo et al. ^2^‘s process included using “drop_outlier_feats” and “drop_outlier_samples” functions from the GitHub repository associated with that work, removing both features and datapoints with absolute values larger than a specified threshold (1e^2^). Applying these to our dataset removed three compounds entirely and 59 well datapoints, potentially resulting in a less difficult classification task for the image-based profile models (documented in the GitHub repository for this work). This removal was not mirrored in the SwinV2, compound structure or CVF models, which predicted the whole dataset shown in Table 2.

### Raw images: Normalization and standardization

Since prior studies typically use processed image-based profiles as the input for either supervised or unsupervised machine learning methods, there is not well-established practice for handling the normalization and standardization of the raw cell painting images. Furthermore, these images are segmented into single cells by CellProfiler before relevant features are measured and extracted. Within this work we use the original images, without segmentation, as input to our computer vision framework. These images are min-max normalized on a channel-wise basis, i.e., each channel is normalized separately according to the formula below, hereafter referred to as per-channel normalization (PCN).

- Min-max normalization:

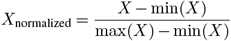

This approach was devised after observing the heterogeneity of pixel intensity values intra-channel within the dataset, suggesting that normalizing across the whole five-channel image would ignore the channel-specific variation which is present in Cell Painted images.

Two approaches, common to the standardization of image-based profiles, were adapted for standardizing the raw images. Within each plate in cpg0016 there are both positive and negative controls as well as untreated wells. The negative controls on the compound plates consist of wells containing cells situated in DMSO solution.

For the positive controls, eight compounds found to have the most distinctly different phenotypic signatures in the CP JUMP1 pilot experiment ^11^, were chosen. These compounds are detailed in Supplemental Table 1 of Chandrasekaran et al. ^16^‘s report, along with their unique identifier in the cpg0016 dataset. They include: Aloxistatin (cysteine protease inhibitor), AMG900 (aurora kinase inhibitor), Dexamethasone (corticosteroid), FK-866 (nicotinamide phosphoribosyltransferase (NMPRTase) inhibitor), LY2109761 (transforming growth factor beta receptor inhibitor), NVSPAK1-1 (p21-activating kinase inhibitor), Quinidine (cinchona alkaloid) and TC-S-7004 (dual-specificity tyrosine phosphorylation-regulated kinase inhibitor). The aurora kinase inhibitor positive control shares its MOA with one of the classes in the kinase inhibitor dataset, as such this compound was excluded when calculating the positive control standardization metrics for this work.

Using the control samples on each plate, the mean, standard deviation, median, and median absolute deviation of the pixel values were extracted from the images of the negative and positive controls samples separately. These statistics were obtained on a channel-wise basis. Two types of standardization could then be applied to the images in the kinase dataset:

- Mean and Standard Deviation:

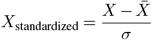
- Median and Median Absolute Deviation (MAD):

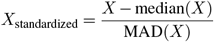

This plate-wise, channel-wise standardization approach is hereafter referred to as PCS. Hyperparameter tuning found median and MAD (MedMAD) PCS to be the more effective, yielding improved model performance. This was likely due to the outliers present in the image pixel values, as both median and MAD are less influenced by extreme values. Tuning also found using the positive controls to be more beneficial than using the DMSO negative control samples. For the CVF model the accuracy using either approach was the same, however using DMSO controls to standardize the image data lead to 2.09% lower F1 score. The impact of PCS and PCN on an example image can be observed in Figure 7. A schematic of the PCS process is included in Supplemental Note S5.

**Figure 7.**
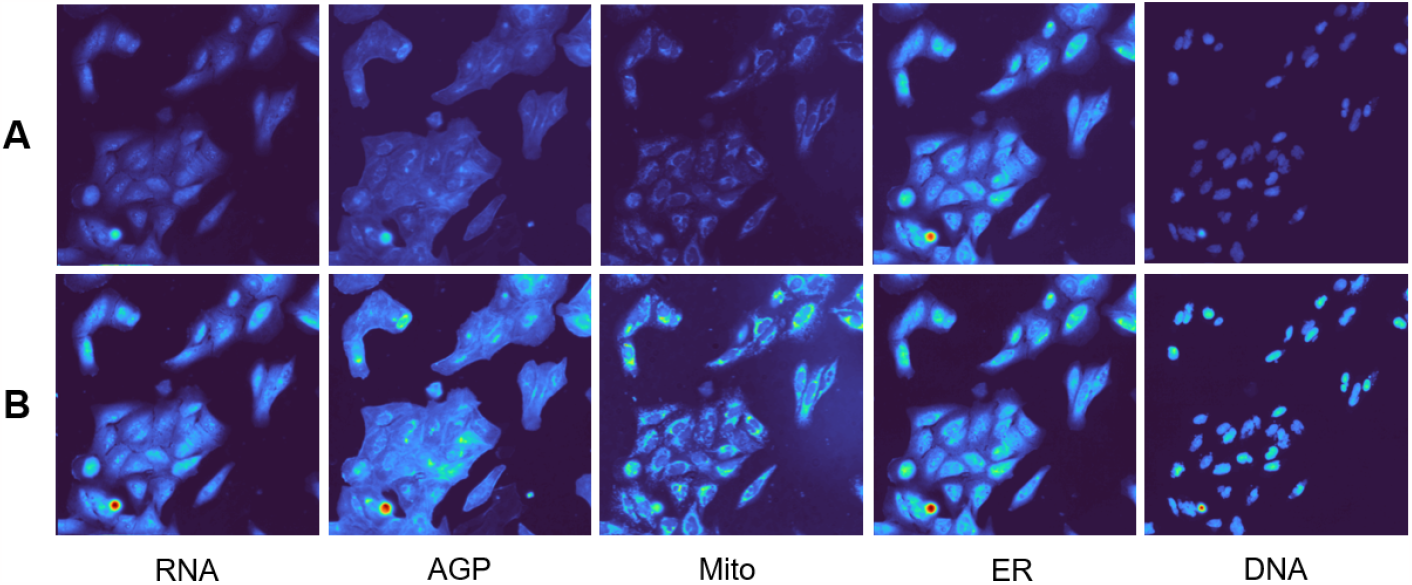
Applying PCN Normalization and PCS Standardization to a Cell Painting Image. A comparison between (A) non-normalized images from one field of view taken across the five cell painting channels (RNA, AGP, Mito, ER and DNA) and (B) the same images after undergoing PCS and PCN. Images are displayed using the “turbo” colormap from the Matplotlib package, to make the individual components more clear.

### Raw images: Data augmentation

Kim et al. ^70^ performed a systematic analysis of common image augmentations for multi-channel microscopy images, finding that a combination of random brightness shifts and intensity changes applied independently to each image channel had the greatest positive impact on model performance. We have extended this approach by adding a probability weighting for each channel, hereby referring to this method as Channel-wise Augmentation (CWA). We found optimal performance occurred when, for each image, the brightness and intensity shift augmentation is triggered with probability P_A_ = 0.8, and for each channel within that image, it is then applied with a secondary probability P_C_ = 0.4. The intensity shift ranged from -0.6 to 0.6, whist the brightness change ranged from -0.1 to 0.1, where both augmentation output images are subsequently clipped in the range [0, 1]. An exploration of the impact of the parameter P_C_’s impact on model performance can be found in Supplemental Note S6.

Additionally, each image was cropped because the images contributed by each source range from 996 × 996-pixels (sources two and five) to 1,280 × 1,080 (source seven). They were cropped to 896 × 896 pixels so the images are divisible into the corresponding patch sizes required by SwinV2. Random vertical and horizontal flips were also applied during training to help combat overfitting.

### Compound Structure

Prior research has shown that incorporating compound chemical structure as a feature can provide a context-free (i.e., cell line agnostic) approach to MOA prediction ^15^. Including this information, alongside cell painting images, lead to a significant improvement in the ability of a multi-layer perceptron (MLP) model to predict MOA.

In this research we have replicated the approach of Tian et al. ^15^ by using the chemical simplified molecular-input line-entry system (SMILES) included within the metadata of the cpg0016 dataset. The SMILES were converted into their molecular structures using the RDKit package function “Chem.MolFromSmiles” before transforming them into 2,048-long Morgan Fingerprint ^71^ vectors using the “GetMorganFingerprintAsBitVect” function of RDKit. We used the Extended-Connectivity Fingerprint (ECFP) ^72^ iteration which encodes the compound’s molecular structure as a binary bitstring representation based on the connectivity of atoms and their neighbouring atoms within a defined radius. The resulting vectorized fingerprint therefore captures the presence or absence of specific substructures or chemical features in a molecule.

### Modelling approaches

For the IBP data, three different machine learning approaches were chosen to test. Random Forest (RF) ^73^ and Support Vector Machines (SVM) ^74^ were trialed due their success in previous MOA or cellular perturbation based tasks. XGBoost (XGB) ^75^ was also selected due to the relative success of tree-based models in prior work, and because of its success in predicting drug-induced cell viability in research using L1000 gene expression profiles ^76^, representations which are similar to the phenotypic profiles in this work. Both RF and SVM were imported from the sklearn package, while XGB was implemented using the dmlc XGBoost implementation.

Additionally, a Multi-layer Perceptron (MLP) model was utilized, based on the results of Way et al. ^3^’s work where it outperformed all other models tested for predicting the MOA of compounds represented by both Cell Painting and L1000 data. The model was adapted from Way et al. ^3^’s original Tensorflow code into Pytorch to be consistent with the other models in this work, but the core elements were retained, which included: six fully connected layers with batch normalization, drop out in the first two layers, combinations of the “ELU” (first, third), “ReLU” (second, third), and “SELU” (fifth layer) activation functions and the Adam optimizer ^77^. For the raw image data, a transformer-based model ^78^ was desired because we aimed to process the whole images directly and required an architecture which would be capable of modelling long-term dependencies in the data, capturing the overall inter and intra-cellular context of the phenotypic changes caused by the kinase inhibitors. The Swin Transformer ^35^ was selected due to its usage of shifting image patch windows between self-attention layers, limiting the computational complexity while still computing attention globally. This was important due to the scaling nature of the number of images for each compound in the dataset (shown in Table 2), whereby 96 unique compounds equate to 5,191 underlying five-channel images. The SwinV2 implementation was chosen due to its ability to train with greater stability given higher resolution input data. This represents one of the first instances of a deep learning, transformer-based model being applied to Cell Painting data for MOA prediction.

EfficientNet ^79^ was used as a comparison for SwinV2, given its popularity in prior cellular perturbation research ^7,15,66,80,81^. EfficientNet is a family of convolutional neural network (CNN) architectures that aim to achieve higher accuracy and efficiency by balancing the model’s depth, width, and resolution by using a compound coefficient to scale each component uniformly. Depth refers to the number of layers in the network, width represents the width of the layers (number of channels), and resolution indicates the input image size. This makes the architecture well-suited to handle the five-channel Cell Painting images. The EfficientNet-B1 implementation was applied in this paper and an extra processing step was added, to resize each input image to 240 × 240 pixels during pre-processing, similar to prior work.

Tian et al. ^15^ showed that incorporating a model trained to classify MOA based on the underlying compound chemical structure can provide a context-free (i.e., cell line agnostic) approach to prediction. Furthermore, by concatenating the output from their structural model with a separate model trained on five-channel Cell Painting data, their overall model performance increased from “a macro-averaged F1 score of 0.58 when training on only the structural data, [to] 0.81 when training on only the image data, and 0.92 when training on both together”. Following the approach of Tian et al. ^15^‘s research, a similar MLP model was constructed to classify data based on the underlying compound’s chemical structure.

### Cell-Vision Fusion (CVF) approach

Once the best performing model architecture and hyperparameters were established for each data modality, the CVF architecture was constructed which incorporated all three modalities, see Figure 2 for a schematic of the approach.

Since there are multiple fields of view for each experimental well, there are many more images than profiles, as each profile is summarized at the well level. Similarly, as there are replicates of each compound spread across different wells, there are many more well-level profiles than there are individual compounds (see Data Selection for full details). Therefore, to prevent the fusion model for overfitting, weight decay regularization of 1e-2 was applied using the AdamW optimizer ^82^, in addition to a dropout of 0.2 for the IBP MLP and 0.5 for the compound structure MLP model. Several different methods of fusion were tested when trying to combine the model outputs, including using a learnable fusion layer, an attention-based combination and concatenation. The most successful method, however, utilized Multimodal Outer Arithmetic Block (MOAB) fusion ^23^, combining the three modalities through four separate arithmetic operations which are concatenated before applying a convolutional layer to the resulting matrix. Min-max normalizing the IBP data resulted in greater performance compared to other normalization approaches when all modalities were combined, likely due to the MOAB combination of different model outputs.

### Analysis and evaluation

XGBoost was selected as the method to investigate the impact of feature selection on model performance as tree-based models are generally not affected by a lack of normalization/ feature scaling. This allowed us to first isolate the impact of feature selection on model performance, before applying normalization and standardization methods. Tree-based models also have a hierarchical structure, which generally makes them more interpretable than deep-learning methods, aligning well with the feature contributions of Shapley values. Furthermore, for generating Shapley values, tree-based methods lead to a more efficient value calculation and greater stability and accuracy of feature value predictions ^83^.

We applied a 5-fold double-stratified cross-validation strategy, similar to Way et al. ^3^, whereby compounds were stratified by MOA, while ensuring replicates of the same compound were retained in the same training, validation or test set. We utilized a training, validation, test split of 70:10:20, ensuring there was at least one compound of each MOA across each test set. Validation data was used to search for optimal hyperparameters before models were trained with the training and validation data, before being evaluated on the held-out test data of each fold.

Prior MOA research has typically used a combination of accuracy, macro-averaged F1 scores, recall and precision to assess model performance ^3,13,15,21^. The area under the precision-recall curve (AUPR) has also been used to represent the trade-off between precision and recall and can be a less optimistic measure than the area under the receiver operating characteristic curve when dealing with datasets with imbalanced, but equally important classes. These metrics were calculated in this paper using the sklearn package.

For the IBP data, predictions are made by the models at a well-level. When calculating model metrics, these class prediction probabilities are averaged across wells (i.e., technical/ experimental replicates) to result in a compound-level prediction. For the image data, this amalgamation has one further step, as predictions are first averaged across the fields within a well, before being combined into average well-level predictions and finally a class prediction for each compound. The structural model is trained at the compound level initially, so no further processing is required.

## ACKNOWLEDGEMENTS

This work was supported by the UKRI/BBSRC Collaborative Training Partnership in AI for Drug Discovery, led by Exscientia in partnership with Queen Mary University of London. The authors would like to thank the Broad Institute and the JUMP Consortium for making the Cell Painting data freely available and accessible. We would also like to thanks John Overington for his feedback on the draft manuscript.

## AUTHOR CONTRIBUTIONS

AL and WD conceived the study. WD devised and implemented the bioinformatics, data processing, analysis and evaluation pipelines. AL, GS and IS supervised the project and provided direction and input towards its progression. WD drafted the manuscript. AL, GS and IS edited and gave feedback on the manuscript.

## DATA AND SOURCE CODE AVAILABILITY

Code to replicate the results of this research, as well as instructions for how to access the underlying data, can be found at the following respository: https://github.com/williamdee1/Cell-Vision-Fusion.

