## Supplementary Information for "Cell-Vision Fusion: A Swin Transformer-based Approach to Predicting Kinase Inhibitor Mechanism of Action from Cell Painting Data"

### S1. Shapley importance values per class

Displayed in Figure S1 are the top ten features for predicting each MOA class when using an XGBoost model, ranked according to Shapley importance value.

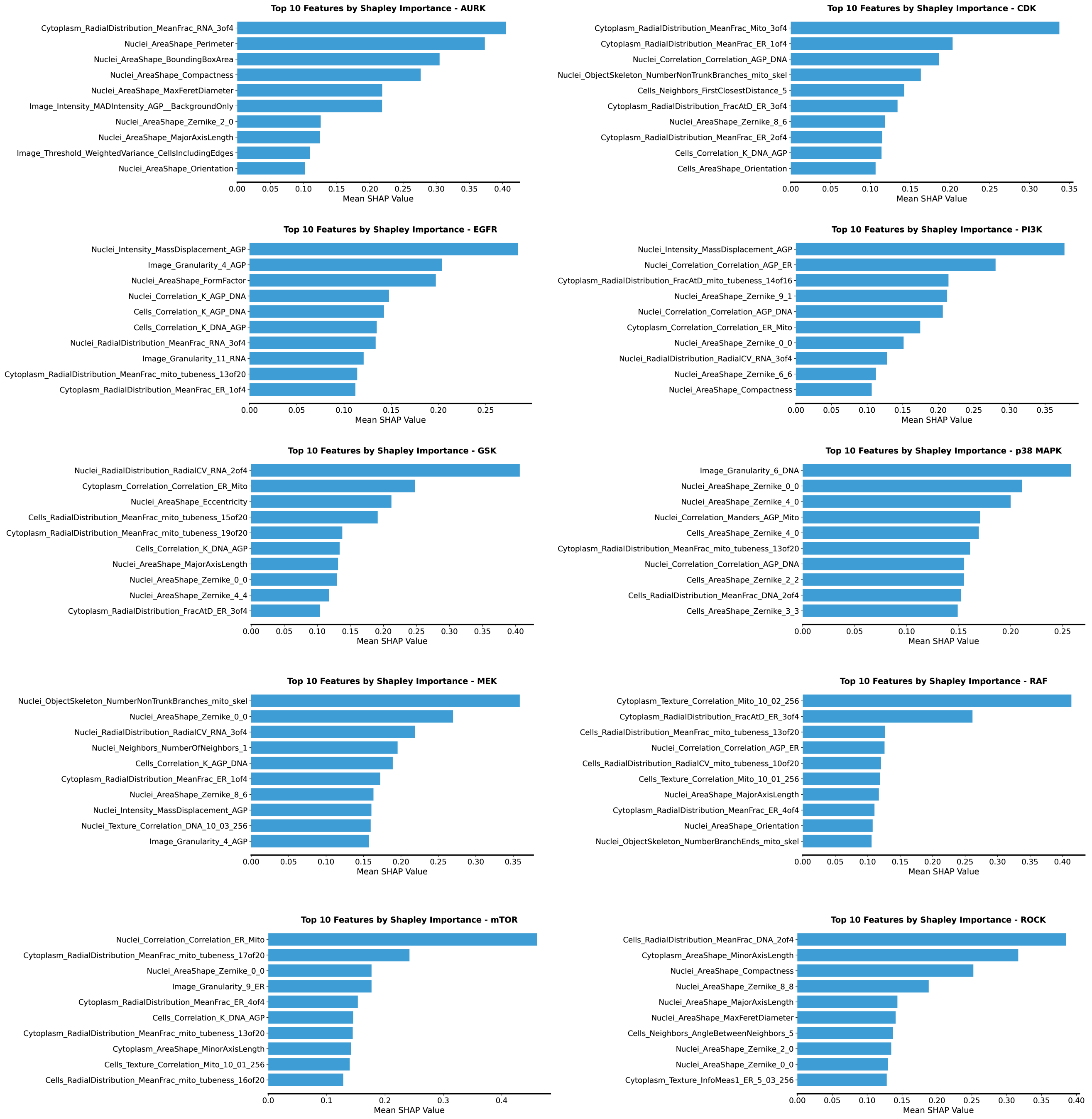


**Supplemental Information, Figure S1.** Shapley importance values for the top ten ranked features for predicting each MOA class in the kinase inhibitor dataset.

### S2. U2OS-specific Kinase inhibitor class selection

#### DepMap gene TPM values

Gene expression TPM values for the U2OS cell line were downloaded from the DepMap Public 23Q2 gene TPM values using the DepMap U2OS ID "ACH-000364". The broad kinase inhibitor classes found in the JUMP cpg0016 data were then associated with the specific underlying genes found in the DepMap data (Figure S2A) before the expression values were summed and sorted at a kinase inhibitor class level (Figure S2B). This provided an idea of the likely expression levels of the protein kinases found in U2OS cells which would then impact how significant an impact inhibiting them would have on the phenotypic response of the cell to each inhibitor.

**A**

```
ki_tgt_dict = {
    'PI3K': 'PIK3CA|PIK3CB|PIK3CD|PIK3CG',
    'EGFR': 'EGFR',
    'p38 MAPK': 'MAPK11|MAPK12|MAPK13|MAPK14',
    'JAK': 'JAK1|JAK2|JAK3|TYK2',
    'RAF': '^ARAF|BRAF|^RAF1',
    'AURK': 'AURKA |AURKB|AURKC',
    'ALK': 'ALK ',
    'SRC': '^SRC |LYN |FYN|YES1|FGR|BLK|HCK|LCK',
    'ROCK': 'ROCK1|ROCK2',
    'MEK': 'MAP2K1|MAP2K2',
    'GSK': 'GSK3A|GSK3B',
    'CDK': 'CDK1 |CDK2 |CDK4 |CDK5 |CDK6 |CDK7 |CDK9 ',
    'VEGFR': 'FLT1|^KDR|FLT4',
    'BCR-ABL': '^ABL1 |^ABL2 |BCR',
    'PDGFR': 'PDGFRA|PDGFRB',
    'FGFR': 'FGFR1 |FGFR2|FGFR3|FGFR4',
    'BTK': '^BTK',    # '^' added to exclude IBTK
    'AKT': 'AKT1 |AKT2|AKT3',
    'mTOR': '^MTOR '
    }
```

**B**

| k_inhib | exp_val |
|---|---|
| MEK | 6.034046 |
| CDK | 6.030705 |
| GSK | 5.909385 |
| AKT | 5.856681 |
| AURK | 5.759314 |
| mTOR | 5.100557 |
| RAF | 5.077632 |
| ROCK | 4.744178 |
| FGFR | 4.057675 |
| p38 MAPK | 3.689417 |
| BCR-ABL | 3.515022 |
| JAK | 3.309181 |
| PDGFR | 3.305077 |
| SRC | 2.444012 |
| EGFR | 2.163499 |
| PI3K | 2.159123 |
| VEGFR | 1.168495 |
| ALK | 0.201634 |
| BTK | 0.124328 |

**Supplemental Information, Figure S2.** A) Shows a dictionary used to map kinase classes identified in the cpg0016 dataset to the genes within the DepMap expression TPM value data. b) displays the value responses for each gene amalgamated at a kinase inhibitor class level, implying that classes that inhibit proteins associated with highly expressed genes (i.e. MEK, CDK inhibitors) will likely show a more significant phenotypic response in U2OS cells compared with those classes where associated gene expression is low (i.e. ALK, BTK).

#### Kinase inhibitor U2OS literature

**Kinase inhibitor classes matched to cpg0016 data:** Phosphoinositide 3-kinase (PI3K), Epidermal growth factor receptor (EGFR), p38 mitogen-activated protein kinase (p38 MAPK), Janus kinase (JAK), Rapidly Accelerated Fibrosarcoma (RAF), Aurora kinase (AURK), Anaplastic Lymphoma Kinase (ALK), Sarcoma kinase (SRC), Rho-associated kinases (ROCK), Mitogen-activated protein kinase kinase (MEK), Glycogen synthase kinase (GSK), Cyclin-dependent kinase (CDK), Vascular endothelial growth factor receptor (VEGFR), Breakpoint cluster region-Abelson kinase fusion protein (Bcr-Abl), Platelet-derived growth factor receptor (PDGFR), Fibroblast growth factor receptor (FGFR), Bruton's tyrosine kinase (BTK), Protein kinase B (AKT) and Mammalian target of rapamycin (mTOR).

From these classes, we considered which ones were appropriate for prediction, given the cpg0016 experiments had been conducted only using the U2OS cell line. Due to the cell type, there were multiple classes which will likely not produce a distinguishable phenotypic response that a model would be able to interpret. The brief rationale for each class is summarized below:

- **Vascular endothelial growth factor receptor (VEGFR):** VEGF action is primarily targeted to endothelial, rather than epithelial cells [(1)]. As such, we would not expect to see a significant phenotypic response from this class of inhibitors when administered in U2OS cells which have an epithelial morphology.
- **The Bcr-Abl fusion protein and Bruton's tyrosine kinase (BTK):** Both are primarily activated/expressed in hematopoietic cells [(2,3)].
- **Platelet-derived growth factor receptor (PDGFR) and Fibroblast growth factor receptor (FGFR) Inhibitors:** PDGFR and FGFR are primarily expressed in mesenchymal cells, including fibroblasts, smooth muscle cells, and pericytes, but not in epithelial cells [(4)].
- **Janus Kinase (JAK):** The JAK-STAT pathway is triggered by the binding of cytokines to their respective receptors. Unlike immune cells, epithelial cells typically do not express high levels of cytokine receptors. Furthermore, JAK inhibitors, especially earlier generations, are known to exhibit low selectivity, leading to wide-ranging off-target effects [(5)]. Due to this, including the JAK class may introduce a lot of noise into the dataset, making it more difficult to classify other, more selective, classes. Despite this, Janus kinases play a central role in inflammation and therefore their impact can be significant across different cell types.

Whilst PI3K and EGFR demonstrated relatively low gene expression values (Figure S2B), the literature review highlighted the importance of the pathways they are directly involved in, as well as the fact that the upstream nature of these two classes of inhibitors should mean that they demonstrate phenotypic responses which will be appropriate to model.

### S3. Image file formats

Table S1 contains a summary of the image file formats for each contributing source to the cpg0016 dataset. The information was extracted from image files downloaded from the cpg0016 AWS s3 bucket.

| | Technical Specifications | | | | | | |
|---|---|---|---|---|---|---|---|
| **Measure** | Source 1, 3, 4, 9 | Source 2, 5 | Source 6 | Source 7 | Source 8 | Source 10 | Source 11 |
| **Resolution** | 1,080 x 1,080 | 996 x 996 | 970 x 970 | 1,280 x 1,080 | 1,024 x 1,024 | 1,000 x 1,000 | 1,080 x 1,080 |
| **Bit Depth** | 16 | | | | | | |
| **Compression** | LZW | Uncompressed | | | | | LZW |
| **Microscope** | Opera Phenix | CV8000 | CV8000 | CV7000 | ImageXpress | CV8000 | Operetta |
| **Resolution Unit** | 3 | 2 | 2 | 2 | n/a | 2 | 3 |

**Table S1.** Technical specifications of images produced by the different cpg0016 data sources, including image resolution, bit depth, compression, and resolution unit.

### S4. Quality control

Columns from the CellProfiler IBP feature sets were extracted and associated to three different quality control measures - blur, saturation and focus. In total, 25 blur features, 10 saturation features and 25 focus features were identified. These features were min-maxed normalized and plotted to show their distribution throughout our kinase inhibitor dataset. Figure S3 shows the distribution of the blur scores, coloured according to the source where the datapoint was produced.

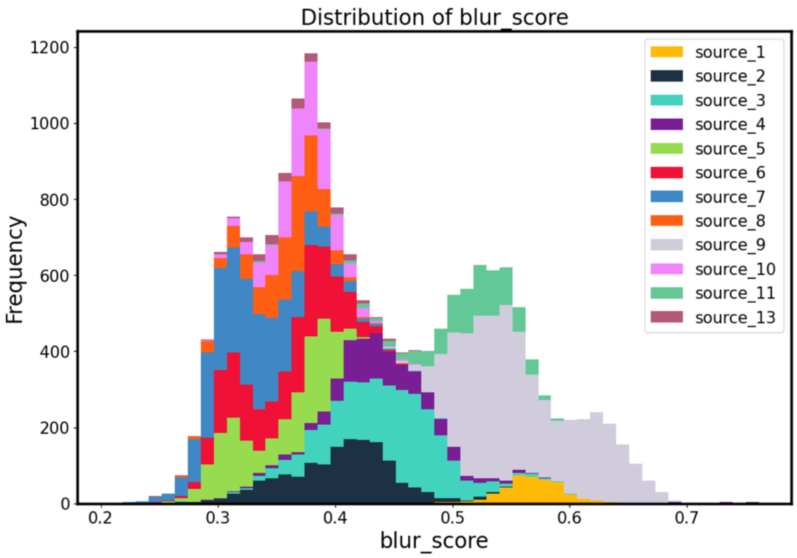


**Supplemental Information, Figure S3.** A plot of the composite, normalized blur scores for each datapoint within the kinase inhibitor dataset, coloured according to which source produced the data. Blur scores were calculated as a mean score from all CellProfiler features associated with the image quality control metric blur, being "ImageQuality_Correlation", "ImageQuality_PowerLogLogSlope" obtained from the CellProfiler manual.

The saturation and focus scores had to be transformed to be comparable to the blur scores. This was due to the fact that the saturation scores were highly skewed, whilst the focus scores were calculated such that a lower score meant lower quality (i.e. low focus), compared to blur scores where a low score meant higher quality (i.e. low blur). Therefore, the saturation scores were transformed into a normal distribution using sklearn's QuantileTransformer module, whilst the focus scores were inverted (Figure S4). On inspection of the images, however, both highly and lowly saturated images were recognised to be poor quality. To combat this, a tanh transform was applied to the saturation scores to ensure all images scoring highly in this metric were reflective of a low quality image (Figure S5).

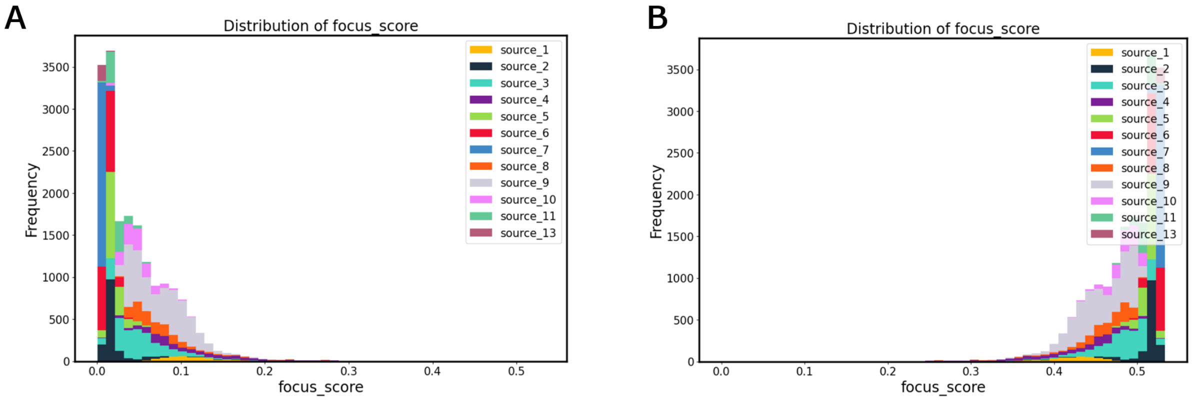


**Supplemental Information, Figure S4.** A) shows the composite mean focus scores for each datapoint before the data was inverted. B) shows the data post-inversion, where a high focus score now equates to a low quality (low focus) image. This transform was performed so the same cutoff could be applied to all three metrics to exclude low quality images. The CellProfiler focus measures used were "ImageQuality_Focus", "ImageQuality_LocalFocus".

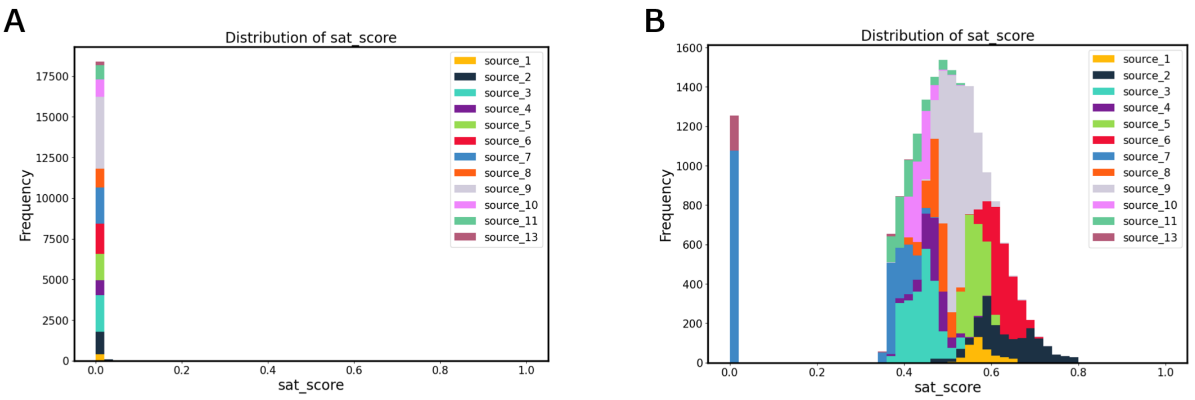


**Supplemental Information, Figure S5.** A) shows the composite mean saturation scores for each datapoint before the data was inverted. B) shows the data after transformation to a normal distribution, followed by a tanh transformation, where a high saturation score now equates to a low quality (lowly or highly saturated) image. This transform was performed so the same cutoff could be applied to all three metrics to exclude low quality images. The CellProfiler saturation measures used were "ImageQuality_PercentMaximal", "ImageQuality_PercentMinimal".

After transforming these measures, they were used to exclude images from selection within our dataset. We applied a 90th percentile cutoff to each metric, ensuring no datapoints, where images which ranked above this threshold for any of the metrics, were included. Figure S6 shows examples of images which were excluded based on this criteria.

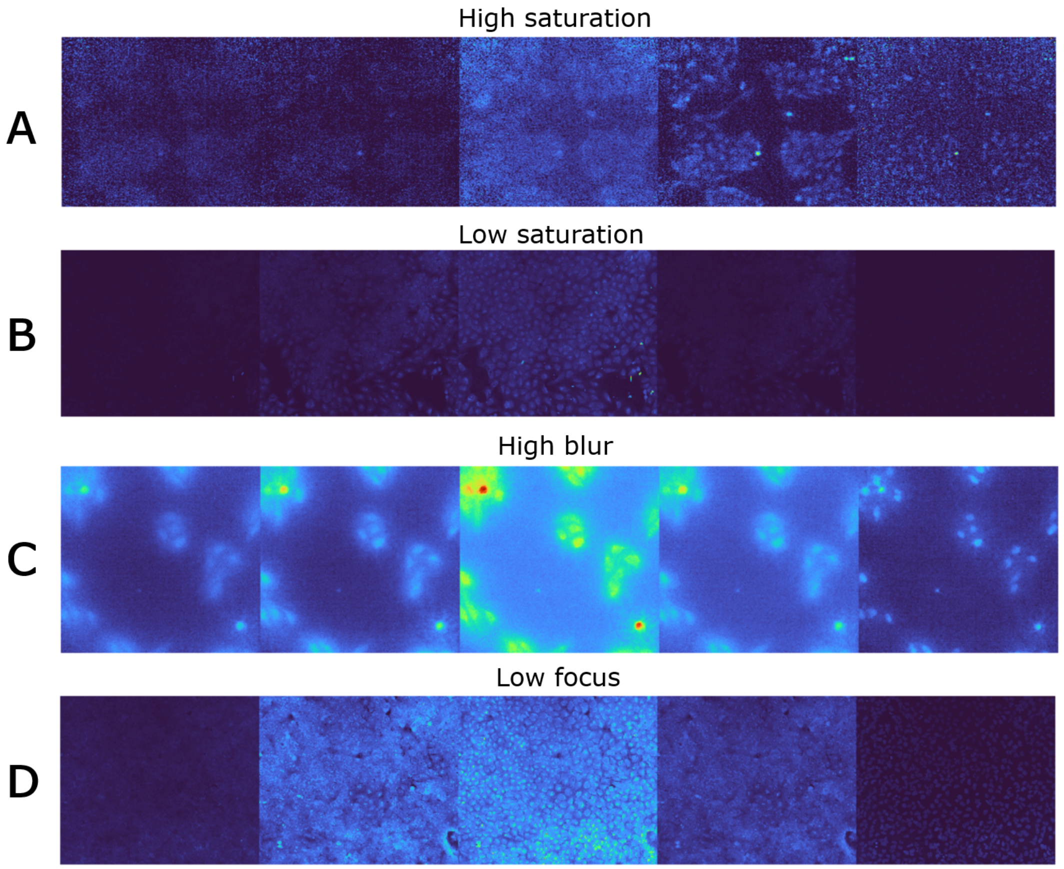


**Supplemental Information, Figure S6.** Example five channel Cell Painting images of datapoints which were excluded based on the quality control metrics defined above, showing examples of: A) high saturation B) low saturation C) high blur and D) low focus. Images are plotted using the 'turbo' colourmap to make the components more clearly visible.

### S5. Plate-wise, channel-wise standardization overview

Figure S7 shows an overview of how PCS is performed on the cpg0016 data to extract the standardization metrics, either median and median absolute deviation or mean and standard deivation, from the positive or negative control samples on each plate.

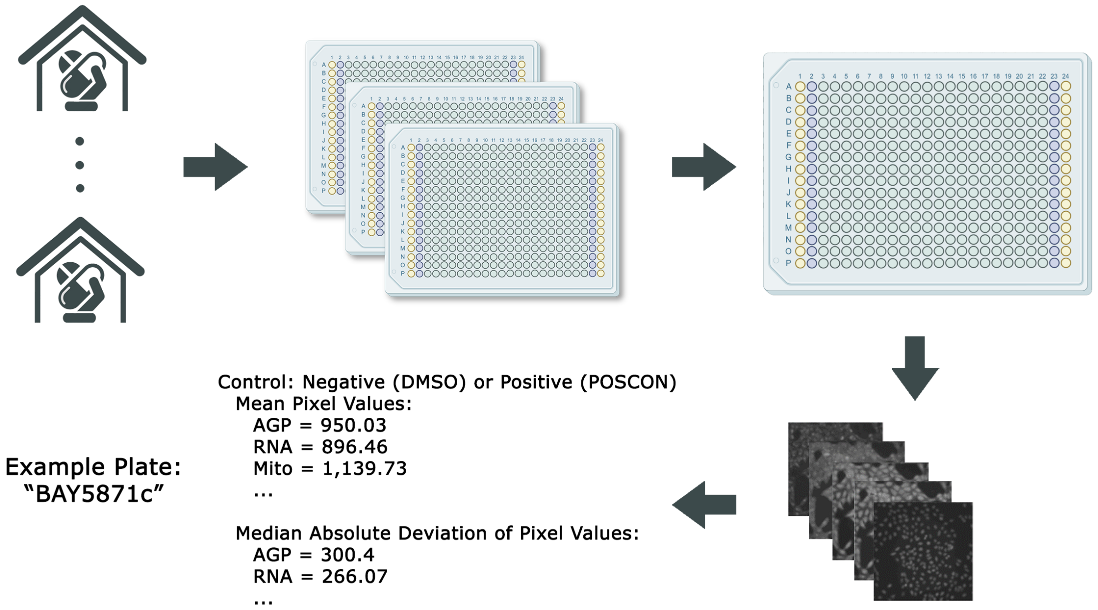


**Supplemental Information, Figure S7.** Overview of the PCS process. Images of either the positive or negative (DMSO) control samples are downloaded from each experimental plate, and the median, mean, median absolute deviation and standard deviation of pixel values are extracted for each channel. These are then aggregated to a plate-level representation of the control response and then these representations can be used to standardize the data from the experimental wells.

### S6. Fusion performance compared to compound clinical phase

The Drug Repurposing Hub and ChEMBL data contain clinical phase information, reflecting how far along in the clinical development pipeline each compound was at the point of inclusion in the dataset. In Figure S8 we compare how this clinical phase impacts model performance, noting that generally as compounds get further along in development they are more accurately identified by the CVF Fusion model. This likely reflects the fact that drugs which have been launched are most likely to effectively impact their intended target with less off-target effects compared to compounds which are still at the preclinical stage.

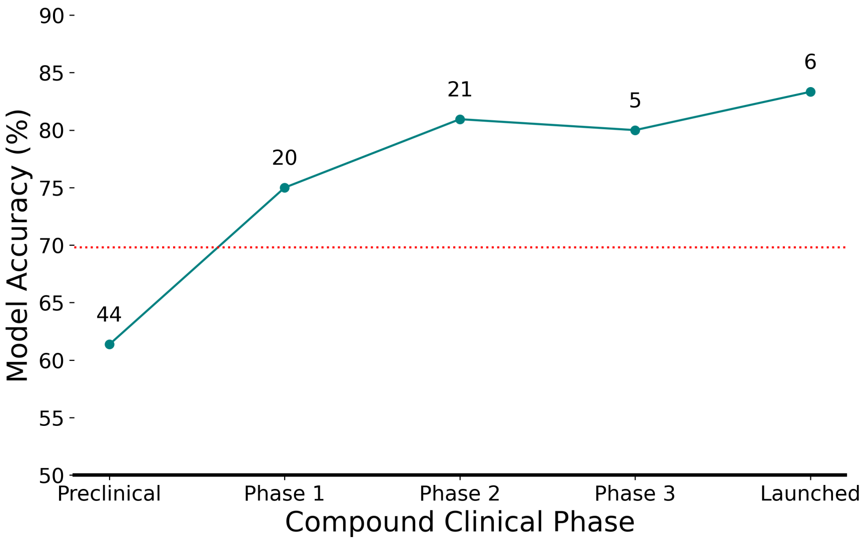


**Supplemental Information, Figure S8.** CVF model accuracy is plotted for compounds grouped according to their clinical phase of development. The points are labelled with a number, reflecting the amound of compounds in that phase in the dataset, i.e., there are 44 in preclinical development out of 96 total in our data. The red dotted line denotes the overall CVF performance achieved, showing that for all compounds at Phase 1 or later, the CVF model classifies them with greater than average performance.

### S7. Study workflow

Figure S9 includes an overview of the workflow used in this study. Initially the JUMP dataset metadata is combined, before MOA labels are assigned to matched compounds using data from both the Drug Repurposing Hub and ChEMBL. Compounds labelled as kinase inhibitors were downloaded before a literature review excluded inhibitors which would likely elicit little to no consistent phenotypic effect in U2OS cells. Quality control was then devised to exclude samples where low quality images were taken by the microscope, this was done using three quality factors - blur, saturation and focus. The remaining samples were included within our kinase inhibitor dataset.

Different approaches to data processing were taken for each data modality, detailed in Figure S9 under the individual modalities. The image-based profiles represent the profiles output by the CellProfiler software which are included with the raw images in the cpg0016 JUMP dataset s3 bucket. Different models were tested for each modality before the best-performing processing techniques and model architectures were combined to form the CVF Fusion model architecture.

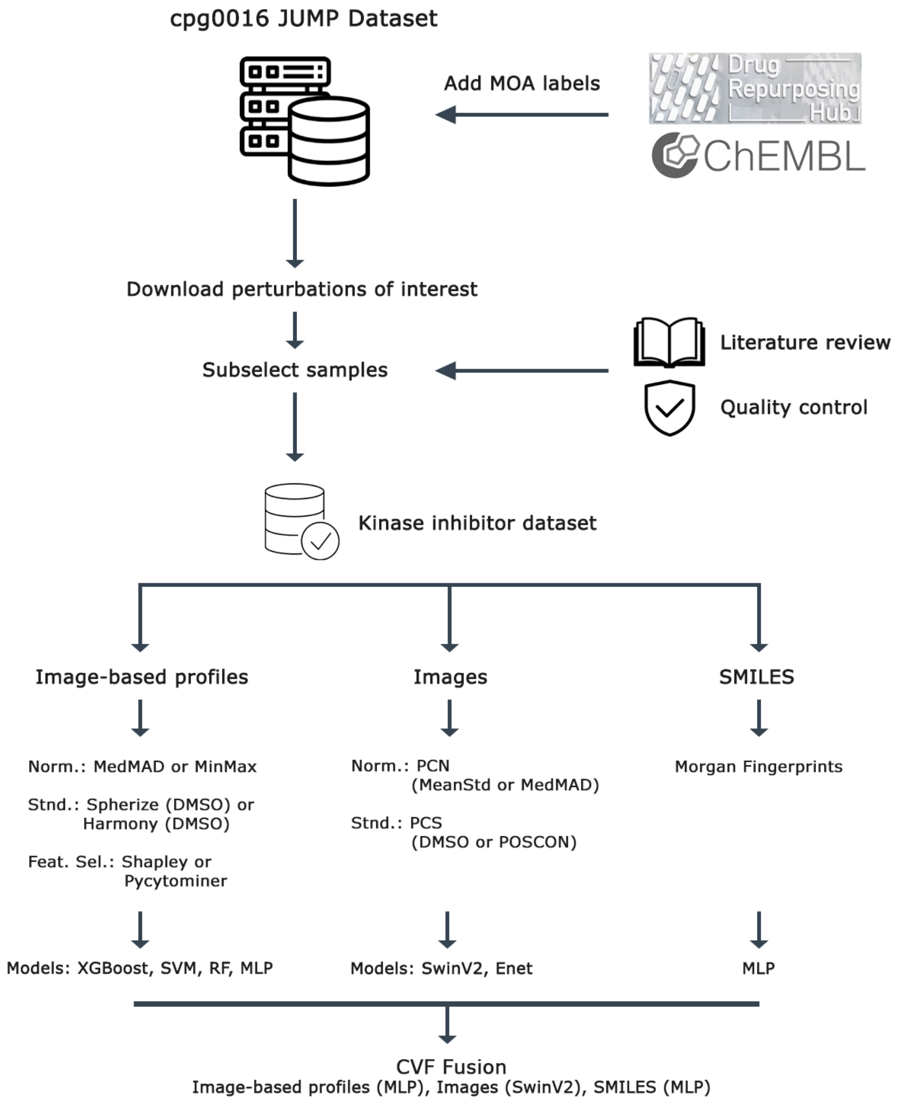


**Supplemental Information, Figure S9.** An overview of the workflow used in this study, describing the dataset selection process, subsequent different processing and modelling approaches, and eventual combination of the optimal approaches into the CVF Fusion architecture.
